# Duplication and Sub/neofunctionalization of *malvolio*, an Insect Homolog of *nramp*, in the Subsocial Beetle *Nicrophorus vespilloides*

**DOI:** 10.1101/135590

**Authors:** Elijah C. Mehlferber, Kyle M. Benowitz, Eilleen M. Roy-Zokan, Elizabeth C. McKinney, Christopher B. Cunningham, Allen J. Moore

## Abstract

Gene duplication has long been thought to play a facilitating role in evolution. With growing numbers of sequenced genomes, increasing numbers of duplicate genes are uncovered with unknown functions. Here we examine *malvolio*, a gene involved in heavy metal transport but that also affects behavior in honey bees and *Drosophila*. There is only one copy of malvolio in honey bees and Drosophila despite its different roles. A phylogenetic analysis in insects suggests that *malvolio* has duplicated multiple times in different orders. To test if the two copies might have different functions, we examined expression levels of *malvolio* in brain, fat bodies, Malpighian tubules, midgut, ovaries, testes and thoracic musculature in the beetle *Nicrophorus vespilloides*. We found that *mvl1* was expressed in all tissues, with highest expression in fat bodies and relatively lower expression in testes, Malpighian tubules, and brain, and ovaries. Expression of *mvl2* differed, with significant expression only seen in brain and midgut. Because *malvolio* has been implicated in behavior, and these beetles have highly developed parenting behavior, we next examined expression during different behavioral states including virgin, mating, preparing resources for offspring, feeding offspring and post care. We found differing expression patterns for the two copies, with *mvl1* increasing in expression during resource preparation and feeding offspring, and *mvl2* decreasing in these same states. Given these patterns of expression, we suggest that *malvolio* in *N. vespilloides* has experienced sub/neofunctionalization following its duplication, and is evolving differing and tissue-specific roles in behavior and physiology.

---

The process of gene duplication is one of the primary mechanisms hypothesized to play a role in the evolution of novel phenotypes (Ohno 1970; Innan and Kondrashov 2010; Ditmar and Liberles 2010; Wagner 2011). When duplicate genes are maintained, one copy often becomes free to mutate and acquire new functions, as it is no longer constrained by the selective pressure to perform its previous role (Ohno 1970; Maere and Van de Peer 2010). This process can take two non-exclusive paths: subfunctionalization or neofunctionalization (Nadeau and Sankoff 1997; Nowak *et al*. 1997; Wagner 1998; Force *et al*. 1999; Maere and Van de Peer 2010). In the former process, both genes lose a portion of their function, so that the two duplicated genes together recapitulate the function of the ancestral gene (Force *et al*. 1999; Maere and Van de Peer 2010). In the latter process, one duplicate evolves a novel function absent from the ancestral gene (Ohno 1970; Maere and Van de Peer 2010). Neofunctionalization may arise following subfunctionalization (Maere and Van de Peer 2010). Given the proposed role that gene duplication has in the production of new phenotypes it follows that more derived organisms with novel traits provide good systems for investigating the divergence of duplicated genes. Genetic influences on behavior may require neofunctionalization of gene duplications to overcome constraints that would otherwise arise through pleiotropy. For example, G-protein coupled receptors are cell surface receptors and have evolved to become diversified and specialized for behavior (Katz and Lillvis 2014). Gene duplication and neofunctionalization has been implicated in the evolution of insect behavior as diverse as *vitellogenin* influences on ant queen and worker social behavior and tasks (Corona *et al*. 2013), olfactory receptors related to shifts to herbivory (Goldman-Huertas *et al*. 2015), and opsin genes related to color vision and foraging preferences (Feuda *et al*. 2016).

With the advent of improved bioinformatics and sequencing, we are acquiring information on genomes of non-model organisms at an accelerated pace, many of which are studied primarily for their novel traits and not for their genetic accessibility. Such genomes provide may reveal duplicated genes that may have previously been unexpected. We recently sequenced, assembled, and annotated the genome of such an organism, the burying beetle *Nicrophorus vespilloides* (Cunningham *et al*. 2015b). Burying beetles (*Nicrophorus spp.*) are unusual among beetles for their parenting behavior. Burying beetles breed on vertebrate carrion, which they shape into a ball, prepare with antimicrobial secretions and bury. The larvae then crawl onto the carcass and one or both parents care for the offspring. Parenting in burying beetles is more than provisioning food for developing offspring, as in many insects, but instead involves direct and extensive prolonged social interactions. Burying beetles not only prepare and maintain a carcass for food, they feed their offspring by regurgitating predigested carrion directly into their mouths. Parenting in this taxon is thus both complex and extensive, and strongly selected as it influences the fitness of offspring (Eggert *et al*. 1998; Lock *et al*. 2004). This behavior is highly derived, and therefore we predicted that sub/neofunctionalization could be important in the evolution of complex parenting. Therefore, we examined this genome for evidence of duplications.

We hypothesized that genes likely to have undergone sub/neofunctionalization in burying beetles are those with a predicted relation to their parenting behavior. Therefore, we focused on genes implicated in the social behavior of other insects. A candidate duplicate gene we found in the genome of *N. vespilloides* is *malvolio*, a heavy metal transporter and homolog of NRAMP family (natural resistance-associated machrophage proteins) in vertebrates (Folwell *et al*.2006). *Malvolio* has been ascribed a role in behavior as well as heavy-metal transport, and is linked to the transition between nurse and forager roles in honey bees (Ben-Shahar *et al*. 2004) and food choice in *Drosophila* (Søvik *et al*. 2017). There is also ample evidence that *Nramp1* (*Slc11a1* in humans) and *Nramp2* (*Slc11a2*) have subfunctionalized in mammals and fish (Techau *et al*. 2007; Neves *et al*. 2011). Other insects such as honey bees and *Drosophila* have only one copy of *malvolio*. It therefore seems likely that this second copy arose in a relatively recent duplication event, and could be unique to beetles.

To test the hypothesis that the duplication of *malvolio* in beetles may facilitate sub/neofunctionalization and effects on behavior, we first examined *malvolio* sequences across insect species and built a gene phylogeny to determine the evolutionary history of duplication in this gene. Next, to ask whether this gene displays behavior consistent with sub/neofunctionalization, we measured gene expression of *mvl1* and *mvl2* across eight tissue types in *N. vespilloides*. *Malvolio* shows evidence of several gene duplications in insects, suggesting something about *malvolio* that lends itself to maintenance after duplication. In *N. vespilloides*, we show that while *mvl1* is expressed ubiquitously and consistently across tissues, *mvl2* only shows expression in the brain and midgut. Finally, we examined expression of the two *malvolio* copies in head tissue collected from beetles before, during and after they were parenting and found changes in expression during parenting in opposite directions for the two copies. We argue that these results provide evidence consistent with the process of sub/neofunctionalization of *mvl*, although it is difficult to tease the two apart. We suggest that the *malvolio* duplicates in *N. vespilloides* are in the process of evolutionary divergence, with neofunctionalization a likely endpoint.

## Materials and Methods

### Biological samples

We maintained *N. vespilloides* as an actively outbred colony at the University of Georgia. We founded the colony with beetles collected from the wild near the University of Exeter, Cornwall, UK and new wild individuals were introduced to the colony yearly to maintain genetic variation. We isolated individuals as larvae and housed them individually in 4×7 cm biodegradable circular deli containers (Eco products, Boulder, CO) filled with 2.5 cm of moist soil (FoxFarm, Samoa, CA). Individuals were kept in an incubator (Percival Scientific, Perry, IA) set at 22 ± 0.1 °C, under a 15:9 light:dark cycle. Upon reaching adulthood they were fed two decapitated mealworms (*Tenebrio*) once a week.

For the comparison of expression across different tissues, we collected eight tissues; brain, fat bodies, hindgut, midgut, thoracic musculature, Malpighian tubules, testes, and ovaries; from 5 virgin female beetles at 26-30 days post adult eclosion (testes came from 5 males of the same age and rearing conditions). These same tissues types were previously examined for octopamine expression in Cunningham *et al*. (2015a), except for testes, but on separate tissue collections. We dissected beetles in ice cold PBS, starting with the brain and then moving on to the internal organs. We cleaned fat and connective tissue from each organ and placed them in separate 1.5 mL vials with 300 µL of RNA later (Applied Biosystems, Foster City, CA) on ice. Dissection times for brains were 10 minutes or less, and the total time of dissections was less than 30 minutes. After dissection, we stored organs overnight in 4°C and then moved them to -20°C until RNA extraction. See Cunningham *et al*. (2015a) for further details.

We collected whole heads from 10 individuals in each of five behavioral states to examine changes associated with changes in behavior: virgins, individuals mated but not provided with the resource necessary to breed, mated individuals provided with a mouse carcass to prepare and that stimulates egg laying, individuals actively caring for and provisioning food to begging offspring, and individuals that had completed parental care and had dispersed away from the carcass and larvae. See Roy-Zokan *et al*. (2015) and Cunningham *et al*. (2016) for further details.

### Identification and comparison of gene sequences

To verify the putative duplication that we found in the genome, we searched for putative *N. vespilloides malvolio* homologs using BLASTp (v2.3.0+; default search settings; Camacho *et al*. 2008) with *Drosophila melanogaster* (NP_524425.2) and *Tribolium castaneum* (XP_967521.1) *mvl* sequences. We obtained sequences from National Center for Biotechnology Information (NCBI) or UniProt. We BLASTed these *mvl* sequences against the proteome produced from the annotated *N. vespilloides* genome (Cunningham *et al*. 2015b). Further identification of putative *mvl1* and *mvl2* of *N. vespilloides* was done by BLASTing (BLASTp, default setting) them into NCBI’s non-redundant insect protein database and by BLASTing (BLASTp, default setting) them into *Drosophila melanogaster* and *Tribolium castaneum* proteomes alone to establish if all of sequences were reciprocal best BLAST’s (RBB) for each other.

All sequences were identified by using the gene and protein databases and the BLAST feature at NCBI (http://www.ncbi.nlm.nih.gov/). To visualize protein conservation across both *mvl* copies, we aligned protein sequences from *N. vespilloides, T. castaneum, H. sapiens*, and *M. musculus* using ClustalW and produced boxshade plots with the Mobyle@Pasteur web portal (http:/mobile.pasteur.fr). For the phylogenetic analysis, we included all insect *mvl* sequences and Mvl proteins available from NCBI, except for *Drosophila spp*. Due to the large number of published Drosophila genomes, and to avoid redundancy, we only included *Drosophila melanogaster*. In order to provide a representative sample of insect species we searched for *mvl* in Lepidoptera but there are currently no assembled and annotated genomes of the order that contain a copy of *mvl*. The NCBI BLAST used both the sequences for Mvl1 (XP_967521.1) and Mvl2 (XP_973779.1) from *T. castaneum*. Protein sequences were then aligned using Clustal Omega (McWilliam *et al*. 2013), and a model test was performed in Mr. Bayes v3.2 (Ronquist *et al*. 2012) to determine the most appropriate model of protein evolution, which was WAG (Whelan and Goldman 2001). A Bayesian phylogenetic analysis was conducted in Mr. Bayes for 5,000,000 generations with a sample frequency of every 100 generations. The consensus tree was compiled after discarding the first 25% of trees sampled and the resultant tree was rooted with human Slc11a1 and SLC11a2, mouse Nramp1 and Nramp2, and *C. gigas* Mvl outgroup.

### Comparison of gene expression

RNA was extracted using a Qiagen RNeasy micro kit (Qiagen, Venlo, Netherlands) for the brain tissue and larval hemolymph and a Qiagen RNeasy lipid kit for all other tissue. The extractions were performed with 350 µL QIAzol (Qiagen) as the lysis buffer and 150 µL chloroform (J.T. Baker, Center Valley, PA). DNA was removed using DNase I (Qiagen) according to manufactures instructions. After the final RNA product was obtained it was quantified with the Qubit 2.0 fluorometer according to manufactures instructions. The RNA was then stored until the time of cDNA production in a freezer set to -80 °C. cDNA was created using 500 ng total RNA and the Quanta Biosciences qScript reverse transcriptase master mix (QuantaBio, Beverly, MA) following the manufactures instructions. The RNA template was then eliminated using RNase H (New England BioLabs, Ipswich, MA) and the single-strand cDNA was quantified using the Qubit 2.0 fluorometer according to manufacturer’s instructions. The resulting cDNA was then stored at -20 °C.

Using the two *malvolio* gene sequences 8 primer pairs (4 primer pairs per gene) were produced by utilizing Integrated DNA Technology (IDT, Coralville, IA, USA) and Primer 3 v. 4.0 (Koressaar and Remm 2007; Untergasser *et al*. 2012). These primer pairs were then validated by estimating PCR efficiency and observing the number of amplicons generated by each pair. The primer efficiency was determined by running a qRT-PCR reaction with stock cDNA (produced using same methods as experimental cDNA from whole body samples) diluted to 1:4, 1:16, 1:64, 1:256 and 1:1024 concentrations, while amplicons were observed in the Melt Curve Analysis. These primer pairs had efficiency levels of 1.805 (*mvl1*) and 1.7852 (*mvl2*).

The quantification of gene expression was accomplished using a qRT-PCR reaction with the Roche LightCycler 480 using Roche LightCycler 480 SYBR I Green Master Mix (Roche Applied Science, Indianapolis, IN, USA). Each biological replicate (N = 5) was run with three technical replicates using 10 µL reactions containing 5 µL SYBR mix, 2 µL of 1.5ng/ µL cDNA, and 3 µL of an equal mixture of forward and reverse primers at 1.33 µmol/L each. The LightCycler was run according to manufacturer’s instructions for the enzyme activation step, followed by 45 cycles of amplification at 60°C and a disassociation curve step to measure the number of amplicons produced in the reaction. Each reaction included the primers (*mvl1*-forward: CGACGATGACGGGAACTTATG reverse: TTGCGATGGATCTGGTGAAG *mvl2*-forward: GGTATCGTGGGAGCAGTTATC reverse: GCTGCTCTCGATGAGGTAATAG *tbp*-forward: CACCCATGACTCCAGCAGAT reverse: ACGTGCATGCAGAGCTATCTT) for *mvl1*, *mvl2*, and TATA binding protein, an endogenous control gene.

### Statistical analysis

Differences in levels of *mvl1* and *mvl2* expression across tissues were quantified using -Δ*C*_T_, or the difference between experimental and control gene expression. Because both versions of *malvolio* were run on the same tissue, same plate, and using the same control gene, the -Δ*C*_T_ values allows us to qualitatively compare expression of the two genes. We made comparisons among the tissues types using ANOVA on the -Δ*C*_T_ values, with specific pairwise comparisons made using Fisher’s Least Significant Difference (LSD) test.

Comparisons of expression of *mvl1* and *mvl2* across different behavioral states were made as described in Roy-Zokan *et al*. (2015) and Cunningham *et al*. (2016), using the -ΔΔ*C*_T_ method with relative expression standardized to virgins. We used virgins as the comparison as this is the behavioral/physiological state of individuals used in the tissue comparison. ANOVA on relative expression was used to determine statistically significant changes in expression.

All data and reagents are available on request. Data will be deposited in Dryad. All accession numbers for sequences used in the phylogenetic comparison are available in supplemental file 2.

## Results

### Phylogenetic analysis of malvolio across insects

Boxshade plots showing sequence homology between *N. vespilloides, T. castaneum, H. sapiens*, and *M. musculus* can be found in supplemental Fig. 1. Phylogenetic analysis (Figure 1) shows that *malvolio* has undergone several independent gene duplications that have been maintained both in insects and other animals as well. Among insects, *malvolio* appears to have duplicated separately in hemipterans, the ancestor of Coleoptera and Diptera, and several times in wasps. Other than wasps, among the Hymenoptera bees and ants have only one copy of *malvolio*. One duplication appears to have been lost in Diptera.

**Figure 1:**
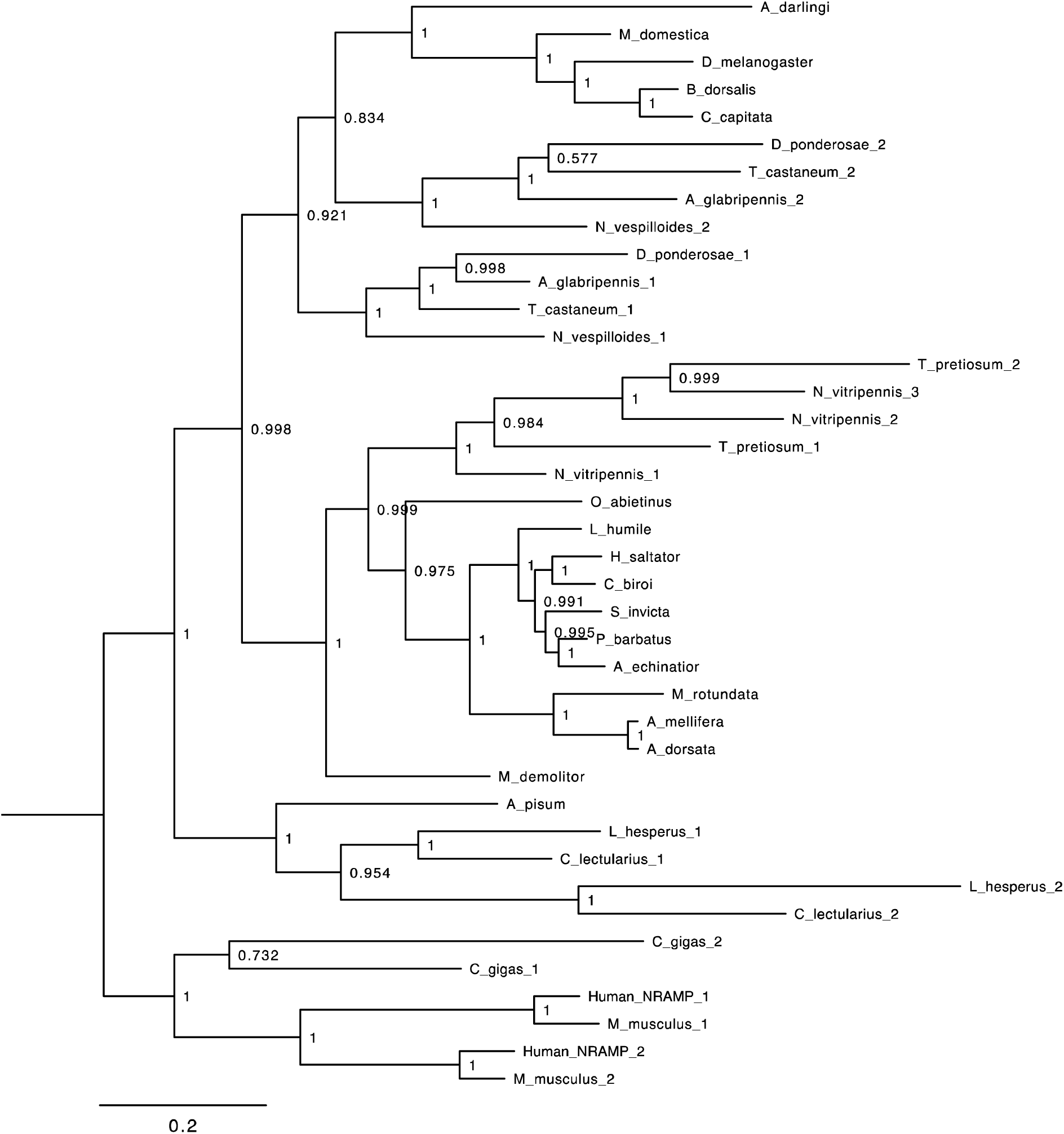
Phylogenetic relationships of *malvolio*. Included in this tree are mammals (human *Homo sapiens*, mouse *Mus musculus*), an oyster (*Crassostera gigas*), a spider (*Latrodectus hesperus*), and several insect orders including Hemiptera (bed bug *Cimex lectularius,* pea aphid *Acyrthosiphon pisum*), Hymenoptera (bees: *Apis dorsata*, *Apis mellifera* and *Metamicroptera rotundata*; wasps: *Orussus abietinus*, *Microplitis demolitor*, *Nasonia vitripennis, Trichogramma pretiosum*; ants: *Acromyrmex echinatior*, *Pogonomyrmex barbatus*, *Solenopsis invicta*, *Ooceraea biroi*, *Harpegnathos saltator*, *Linepithema humile*), Coleoptera (beetles: *Tribolium castaneum*, *Nicrophorus vespilloides*, *Anoplophora glabripennis*, *Dendroctonus ponderosae*) and Diptera (flies: *Ceratitis capitata*, *Bactrocera dorsalis*, *Drosophila melanogaster*, *Musca domestica*, *Anopheles darlingi*).

### Differences in expression in different tissues

Expression of *mvl1* in brain, fat bodies, Malpighian tubules, midgut, ovaries, testes and thoracic musculature varied across the different tissue types (F_7,32_ = 44.361, P < 0.0001). Expression in fat bodies was statistically significantly higher than other tissues (Figure 2a). Hindgut, midgut and thoracic musculature had moderate levels of expression, while expression was relatively low in testes, Malpighian tubules, brains, and ovaries.

**Figure 2:**
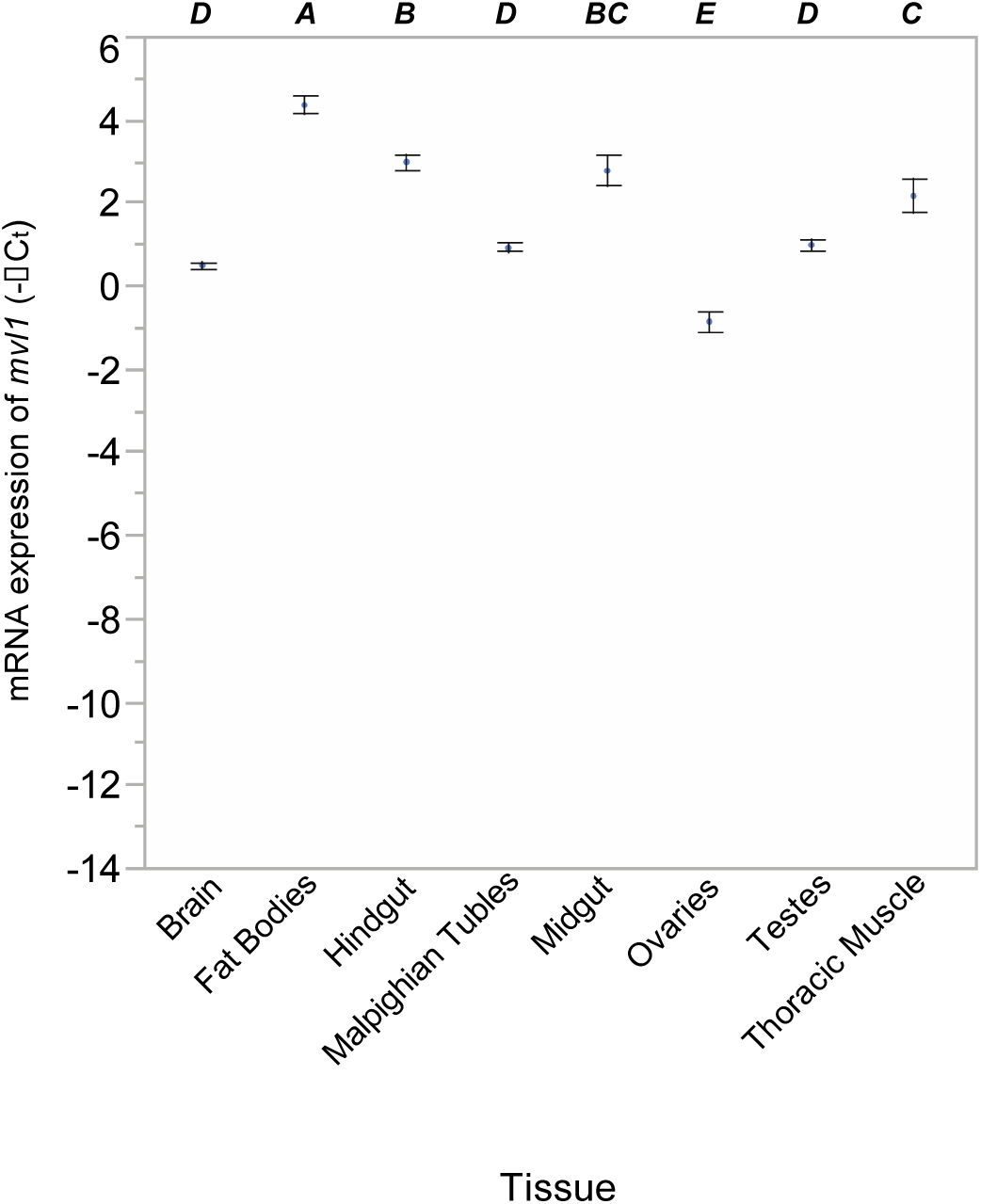
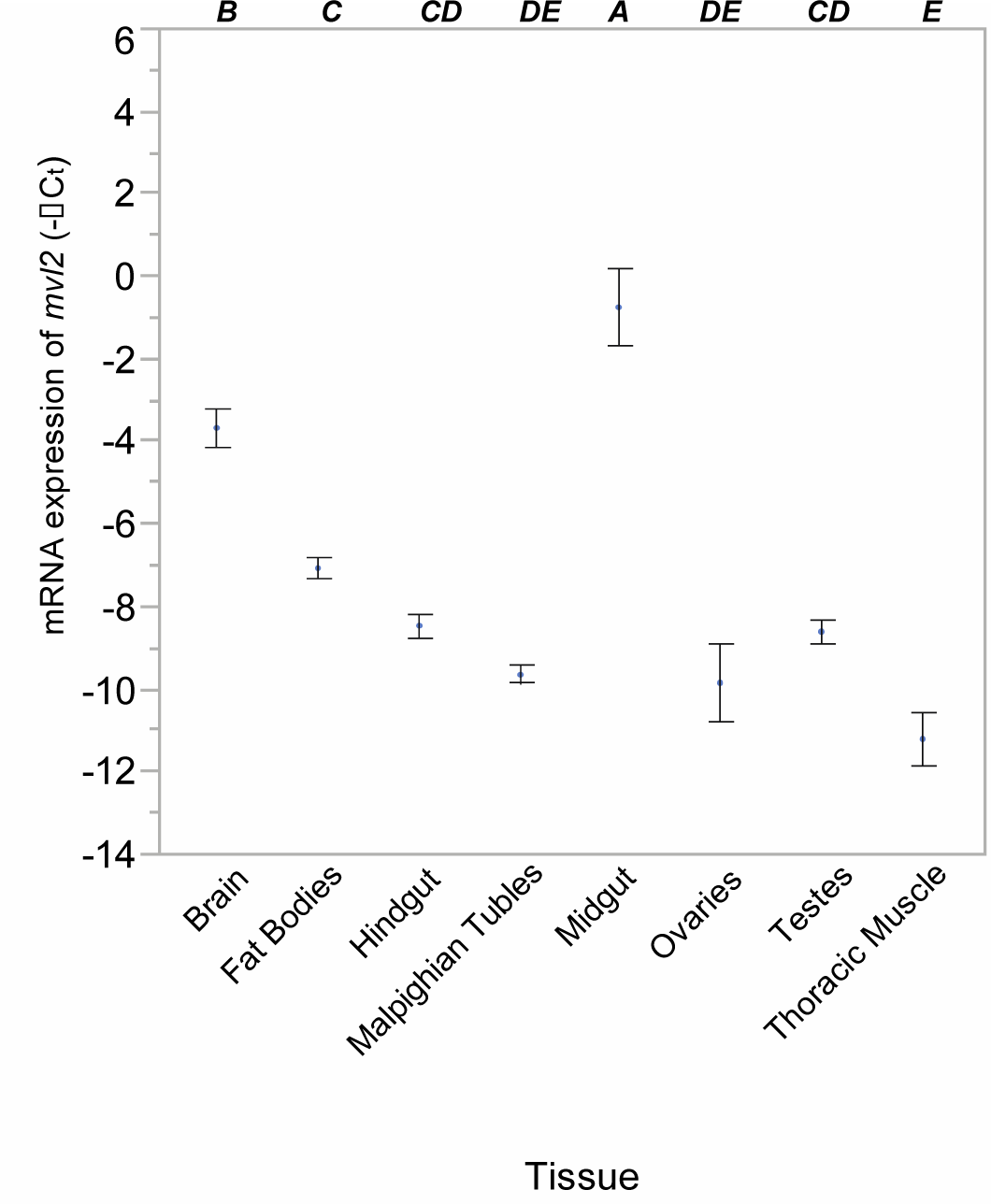
mRNA expression of *mvl1* (2a) and mvl2 (2b) calculated as -Δ*C*_T_. Significant differences in expression are indicated by the letters above each tissue.

**Figure 3:**
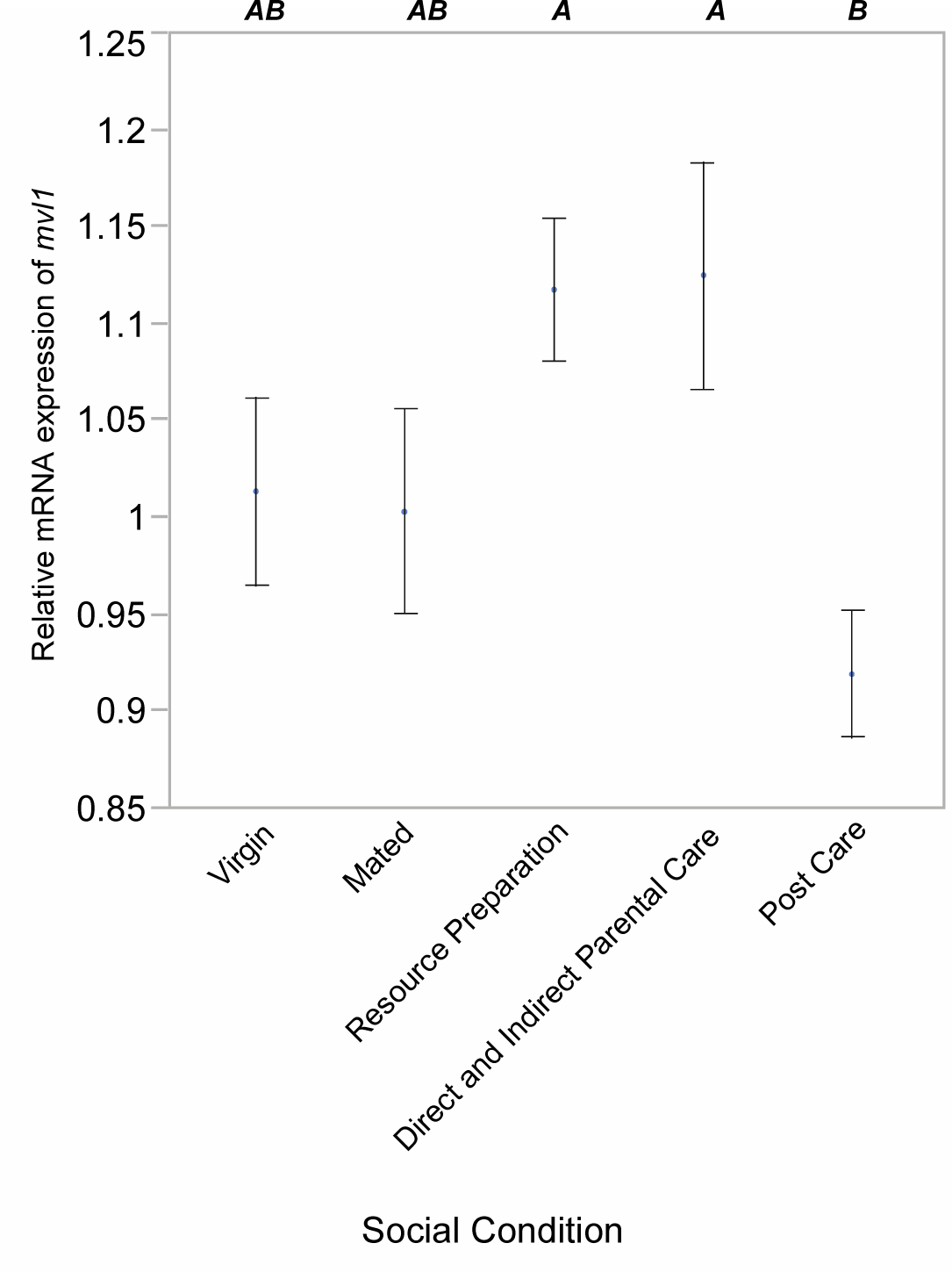
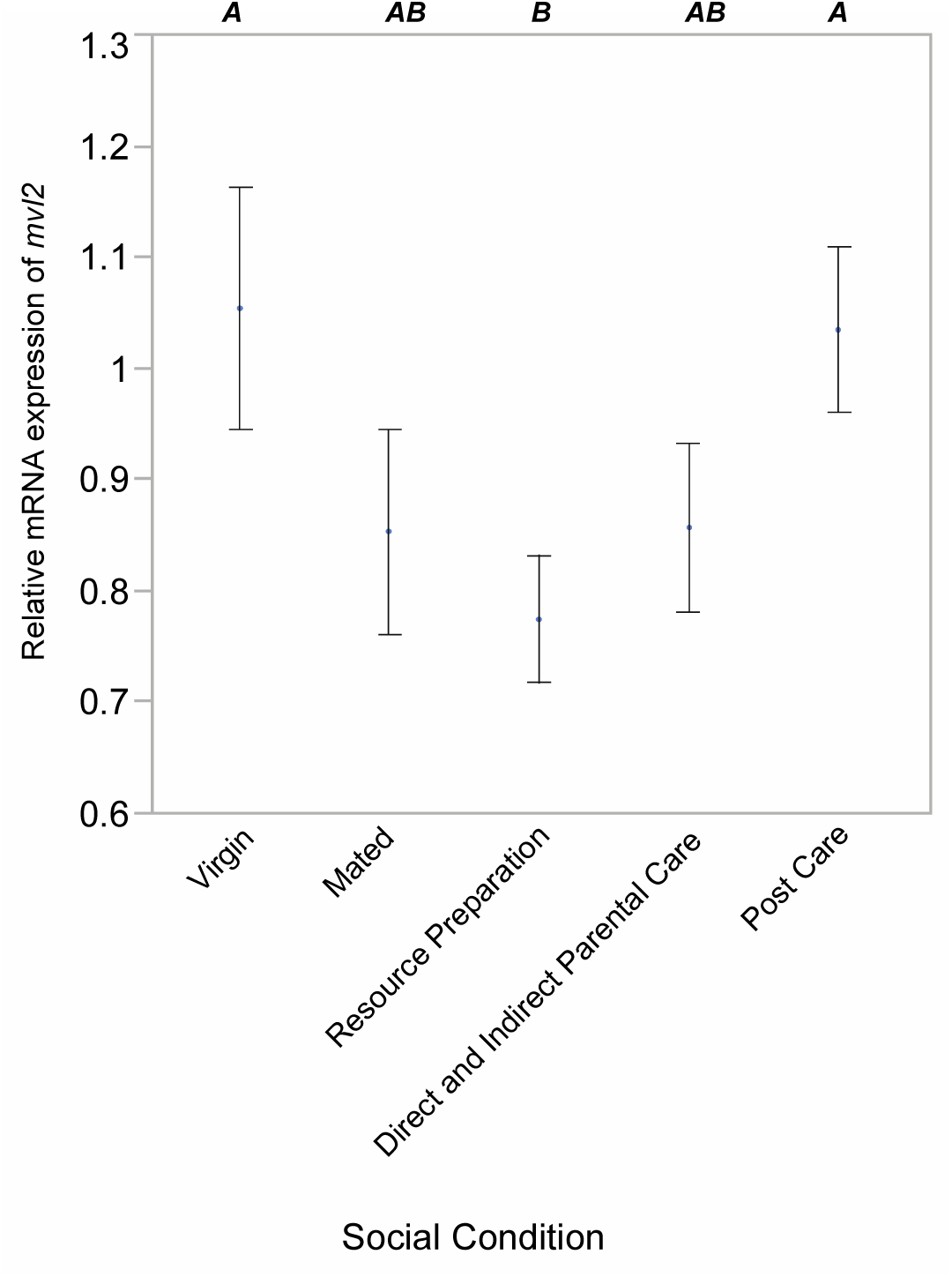
Relative expression of *mvl1* (3a) and *mvl2* (3b) in female heads across different physiological/behavioral states. Virgin females had no social experience or exposure to a carcass resource required for mating and oogenesis; mated females were placed with a male for 48 hours but not provided with a carcass; resource preparation were mated females provided with a carcass for 48 hours to prepare for reproduction and provisioning of offsprig; direct and indirect care were females sampled during the most active period of direct feeding of offspring and in the act of regurgitating food to the offspirng; post care females had dispersed from the carcass and had no further interactions with larvae for at least 24 hours. All individuals were 21 days of age when sampled. Significant differences in expression are indicated by the letters above each state.

Expression patterns across tissues of *mvl2* differed from those of *mvl1* (Figure 2b). Overall, there was statistically significantly different expression across the different tissue types (F_7,32_ = 37.420, P < 0.0001) although in all tissues expression was much lower than that of *mvl1*. Expression was highest in midgut and brain, with low expression in fat bodies, hindgut and testes. Expression was negligible, and sometimes undetectable, in Malpighian tubules, ovaries and thoracic muscle (Figure 2b).

### Differences in expression in heads across different behavioral states

Expression patterns across behavioral states differed for *mvl1* compared to *mvl2*. Overall, there was statistically significantly difference in expression across the behavioral states in *mvl1* (F_4,45_ = 3.4087, P = 0.077) (Fig 4a), with a significant increase in expression in resource preparation (P = 0.0044) and caring for offspring (P = 0.0032). There was no overall statistically significant difference in expression across behavioral states for *mvl2* (F_4,43_ = 2.2682, P = 0.077) (Fig 4b) although there was a strong trend for decreased expression during resource preparation and parental care. In pairwise comparisons, resource preparation showed significantly lower expression than either virgin (P = 0.019) or post care (P = 0.0285). Regardless, the patterns of expression are opposite for *mvl1* and *mvl2* comparing behavioral states.

## Discussion

Gene duplication is a major factor in evolution (Ohno 1970; Innan and Kondrashov 2010; Wagner 2011), particularly where there is neofunctionalization, as the duplicated gene can permit access to variation that may have otherwise been constrained. Here we examined *malvolio*, a gene that typically functions in heavy metal transport. Examining the genome of the subsocial beetle, *N. vespilloides*, we found that *malvolio* was duplicated in this insect. Two other factors suggested that it would be informative to examine this duplication further; first, *malvolio* also plays a role in social behavior in bees (although they have only one copy), and second, *malvolio* is the homolog of *Nramp* in vertebrates, a gene that is duplicated and subfunctionalized (Techau *et al*. 2007; Neves *et al*. 2011). Our study has two components. In our phylogenetic analysis, we found that duplication of this gene was not unique to *N. vespilloides* or insects in general; *malvolio* was often independently duplicated and maintained in multiple insect lineages. In our expression studies, we found that *mvl1* and *mvl2* display both tissue and behavior specific expression patterns. Moreover, level of expression in the brain depended the behavioral state of the insect, with differential expression of both copies of *malvolio*, albeit in opposite direction, associated with parenting behavior.

Contrary to our hypothesis that insects have only one copy of *malvolio*, our phylogenetic analysis showed that many species of insects have duplicate copies and, furthermore, that these duplications appear to have occurred in multiple lineage specific events. *Malvolio* duplicates have arisen and persisted in at least three different insect lineages (and possibly again in wasps, leading at least some to have three copies of *malvolio*). Given the tendency of duplicated genes to remain redundant and eventually be removed from the genome it suggests that *malvolio* may possess qualities that have been found to encourage persistence after a duplication event (Kondrashov *et al*. 2002; Papp *et al*. 2003; Marland *et al*. 2004; Jordan *et al*. 2004; Davis and Petrov 2004).

*Nicrophorus vespilloides* is not a genetic model organism, although we have a sequenced genome (Cunningham *et al*. 2015b). Instead, this species is interesting for its unusually elaborate parenting and social interactions (Parker *et al*. 2015). To examine whether sub/neofunctionalization may be responsible for the maintenance of both *malvolio* duplicates in *N. vespilloides*, we examined tissue-specific expression of both genes. Our data show that *mvl1* is expressed in all eight measured tissues, with relatively low variance in gene expression within a tissue. In contrast to *mvl1*, expression of *mvl2* was limited to only two tissues, the brain and the midgut. This pattern is roughly consistent with tissue and stage-specific data from *Tribolium*, which also shows high and ubiquitous *mvl1* expression as opposed to low and inconsistent, but detectable, *mvl2* expression (Dippel *et al*. 2014). This suggests that *mvl1* may have maintained a conserved homeostatic role throughout the coleopteran lineage, consistent with the necessity of manganese transport on the cellular level (Culotta *et al*. 2005). Differences in expression between specific tissues may be related to other well-established functions of *mvl* and its homologues, such as intercellular immunity (Evans *et al*. 2001; Cellier *et al*. 2007). *mvl2*, on the other hand, is clearly not required for basic tissue function, and thus may be released from pleiotropic constraint.

To further examine the possibility that the function of *mvl2* has diverged from that of *mvl1* in *N. vespilloides*, we examined the expression patterns of both genes in the head in relation to reproductive and parental care behavior. Previous research has shown that genes differentially expressed during parenting are detected in these samples (Parker *et al*. 2015; Roy-Zokan *et al*. 2015; Cunningham *et al*. 2016, 2017). In terms of having a function in behavior, *malvolio* is involved in caste differentiation in honey bees (Ben-Shahar *et al*. 2004) as well as feeding behavior in *Drosophila* (Søvik *et al*. 2017), and therefore represents a strong candidate for influencing social behavior in insects. In particular, we have hypothesized that feeding pathways are co-opted to influence parental provisioning behavior (Cunningham *et al*. 2016, 2017). We found that the two copies do show differences in expression in head tissue associated with changes in behavior and social interactions. Whereas *mvl1* increases expression during parenting, *mvl2* appears to decrease during the same behavioral stages. These opposing expression patterns suggest that even though both gene copies have retained roles in social behavior, these roles have clearly diverged.

Given the tissue and stage-specific expression patterns of *mvl1* and *mvl2*, it appears likely that these genes have undergone either sub-or neofunctionalization in burying beetles. However, it remains unclear which process has occurred. Data from honey bees, in which *malvolio* is not duplicated, shows that a single copy can account for both behavioral and other gene functions (Ben-Shahar *et al*. 2004), suggesting that divergence in gene function between copies could be obtained by subfunctionalization alone. However, it this were the case, we would predict that one copy would have completely lost its association with behavior in *N. vespilloides*. Instead, we observe the evolution of opposing gene expression patterns between copies, meaning the expression patterns of at least one gene copy must be derived. Given the highly divergent and elaborate social interactions during parenting, and extensive parenting, in this species this suggests that *malvolio* may be co-opted for further behavioral evolution. Furthermore, divergence in tissue specific expression patterns, as we observe here, is often associated with neofunctionalization (Huminiecki and Wolfe 2004, Li *et al*. 2005). Therefore, our data is consistent with neofunctionalization. It may be that the expression patterns of *mvl2* in the brain and midgut are still evolving, and understanding whether expression is being gained or lost in these tissues along with explicitly functional studies would help resolve this.

In conclusion, *N. vespilloides* produces two copies of *malvolio*, both of which are expressed, but expression depends on the tissue examined. Duplications of *malvolio* are not unusual among insects, as they appear to have arisen independently and been maintained in several other species of insects. Our expression data suggest that, in *N. vespilloides*, *malvolio* has experienced neo-functionalization following its duplication, with an enhanced role in behavior. Further functional studies are needed to eliminate subfunctionalization but our expression data suggest that the two copies are not equivalent. Finally, we further suggest that the predilection for duplicates of this gene to be maintained may reflect a tendency for sub or neofunctionalization of *mvl* in other systems as well.

## Acknowledgments

We thank Trish Moore and Michelle Ziadie for helpful discussions. This research was supported by a National Science Foundation grant IOS-1326900 to AJM, with ECMe supported by an REU supplement.

